# Morphological deficits of glial cells in a transgenic mouse model for developmental stuttering

**DOI:** 10.1101/2024.01.04.574051

**Authors:** Afuh Adeck, Marissa Millwater, Camryn Bragg, Ruli Zhang, Shahriar SheikhBahaei

**Author notes:** Corresponding author: Shahriar SheikhBahaei, PhD; Neuron-Glia Signaling and Circuits Unit, National Institute of Neurological Disorders and Stroke, National Institutes of Health, Bethesda, MD 20892, USA.

## Abstract

Vocal production involves intricate neural coordination across various brain regions. Stuttering, a common speech disorder, has genetic underpinnings, including mutations in lysosomal-targeting pathway genes. Using a Gnptab-mutant mouse model linked to stuttering, we examined neuron and glial cell morphology in vocal production circuits. Our findings revealed altered astrocyte and microglia processes in these circuits in Gnptab-mutant mice, while control regions remained unaffected. Our results shed light on the potential role of glial cells in stuttering pathophysiology and highlight their relevance in modulating vocal production behaviors.

## 1 INTRODUCTION

Vocal production is a complex motor behavior characterized by the intricate coordination of activities of several brain regions. Although comprehensive brain circuits for speech production have not been fully elucidated, several brain regions, including the cerebral cortices, midbrain, and brainstem regions, have been demonstrated to show substantial involvement in vocal production behaviors (Jürgens, 2002; Simonyan and Fuertinger, 2015; Jarvis, 2019). Speech disorders are prevalent across all cultures, with stuttering being the most common neurodevelopmental speech disorder (Yairi and Ambrose, 2013). Developmental stuttering (also known as child-onset fluency disorder) is defined by involuntary repetitions, prolongations, and frequent pauses or hesitations in speech, known as silent “blocks” (American Psychiatric Association, 2013; Bloodstein et al., 2021).

It is now accepted that genetic substrates play a significant role in the pathophysiology of stuttering (Ambrose et al., 1993; Frigerio-Domingues et al., 2019). Although recent studies have identified specific mutations in several genes with proposed links to developmental stuttering (Kang et al., 2010; Shaw et al., 2021; Polikowsky et al., 2022; Below et al., 2023), few of those mutations have been shown to have a causative effect. Among them, specific mutations in genes necessary for the lysosomal-targeting pathway (i.e., *GNPTAB*, *GNPTG*, *NAGPA*, and *AP4E1*) have been studied in human and animal models (Riaz et al., 2005; Kang et al., 2010; Barnes et al., 2016; Kazemi et al., 2018). We and others have hypothesized that glial cells may play a significant role in the pathophysiology of stuttering (Han et al., 2019; Maguire et al., 2021; Turk et al., 2021; SheikhBahaei et al., 2023b), however, direct evidence to support this hypothesis is lacking, and the extent of the effect of the mutations on glial cells remains unclear.

Here, we used a mouse model for stuttering disorder (*Gnptab*-mutant mouse) that is linked to the disorder in humans (Kang et al., 2010; Barnes et al., 2016) and systematically studied the morphological characteristics of neurons and glia cells (astrocytes, microglia, and oligodendrocytes) in brain regions involved in vocal production and control regions (i.e., regions that are not directly involved in production of vocal behaviors). We found that in the *Gnptab*-mutant mice, morphometric properties of astrocyte and microglia processes are affected in vocal production circuits. Since glia cells possess unique molecular characteristics and roles that are specific to different brain circuits, insights into glial cells in vocal production circuits may offer an advanced understanding of the pathophysiology of stuttering.

## 2 METHODS

### 2.1 Animals

Adult male mice (*Gnptab*-mutant and control littermates, 3-5 months old) were used in this study. All procedures were performed following the *Guide for the Care and Use of Laboratory Animals* and were approved by the Animal Care Use Committee of the Intramural Research Program of the National Institute of Neurological Disorders and Stroke. Homozygous mutant animals were considered *Gnptab*-mutant, and age-matched wildtype littermates were used as control mice. Mice were housed in a temperature-controlled facility with a normal light-dark cycle (12h:12h). Access to food and water was provided *ad libitum*.

### 2.2 Tissue processing and immunohistochemistry

For perfusion-fixation, mice were deeply anesthetized using Isoflurane (5%). When the toe-pinch reflex was undetectable, the mice were perfused transcardially with ice-cold saline solution (0.01M), followed by 4% paraformaldehyde (PFA) fixative as described before (Sheikhbahaei et al., 2018a; Turk and SheikhBahaei, 2022). Briefly, after cryoprotection with 30% sucrose, the brains were sectioned coronally at 50 µm with a freezing microtome. The free-floating tissue sections were incubated at 4°C for 24 h with primary antibody for glial fibrillary acidic protein (GFAP), ionized calcium-binding adaptor molecule 1 (Iba1), anti-proteolipid protein (PLP), or neuronal nuclear protein (NeuN)/Microtubule-associated protein 2 (MAP2), to label astrocytes, microglia, oligodendrocytes, and neurons, respectively (Table 1). Subsequently, the sections were incubated with specific secondary antibodies conjugated to the fluorescent probes (Table 2) for 1.5 h at room temperature. A confocal laser scanning microscope (Zeiss LSM 510) was used to obtain low or high-magnification (20X or 40X, respectively) images from cortical and midbrain regions of interest (Figure 1). We surveyed and imaged astrocytes from each region spanning the rostral-caudal axis for cortical and striatal astrocytes. To minimize variations in processing tissues, one investigator sectioned all the brains, and all the tissue sections were immunostained with identical solutions and processed by the same investigator.

**Figure 1.**
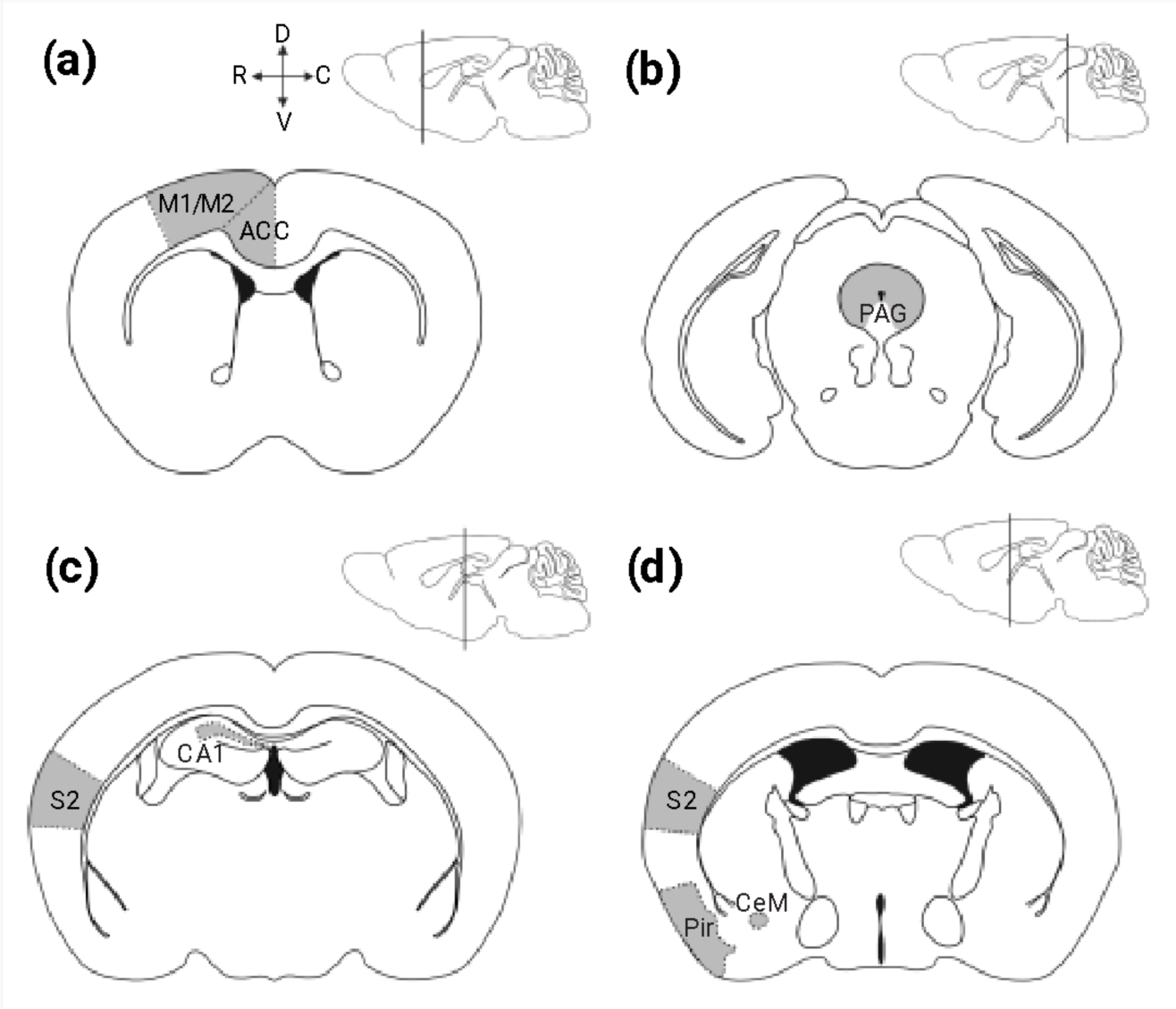
Schematic drawing of adult mouse brain depicting regions of interest for vocal production. (**a-d**) Sagittal views illustrate the regions of interest moving rostral to caudal, with the vertical lines indicating the location of the regions of interest (light gray) in coronal slices. (**a**) Illustration of a coronal section containing primary motor cortex (M1), secondary motor cortex (M2), and anterior cingulate cortex (ACC). (**b**) Schematic drawing of the central nucleus of the amygdala (CeM), secondary somatosensory cortex (S2), and piriform cortex (Pir). **c**) Illustration of hippocampus CA1 region in a coronal slice. **d**) Location of the periaqueductal gray (PAG) is illustrated in a coronal section from the midbrain. d, dorsal; m, medial; l, lateral; v, ventral.

**Table 1.**
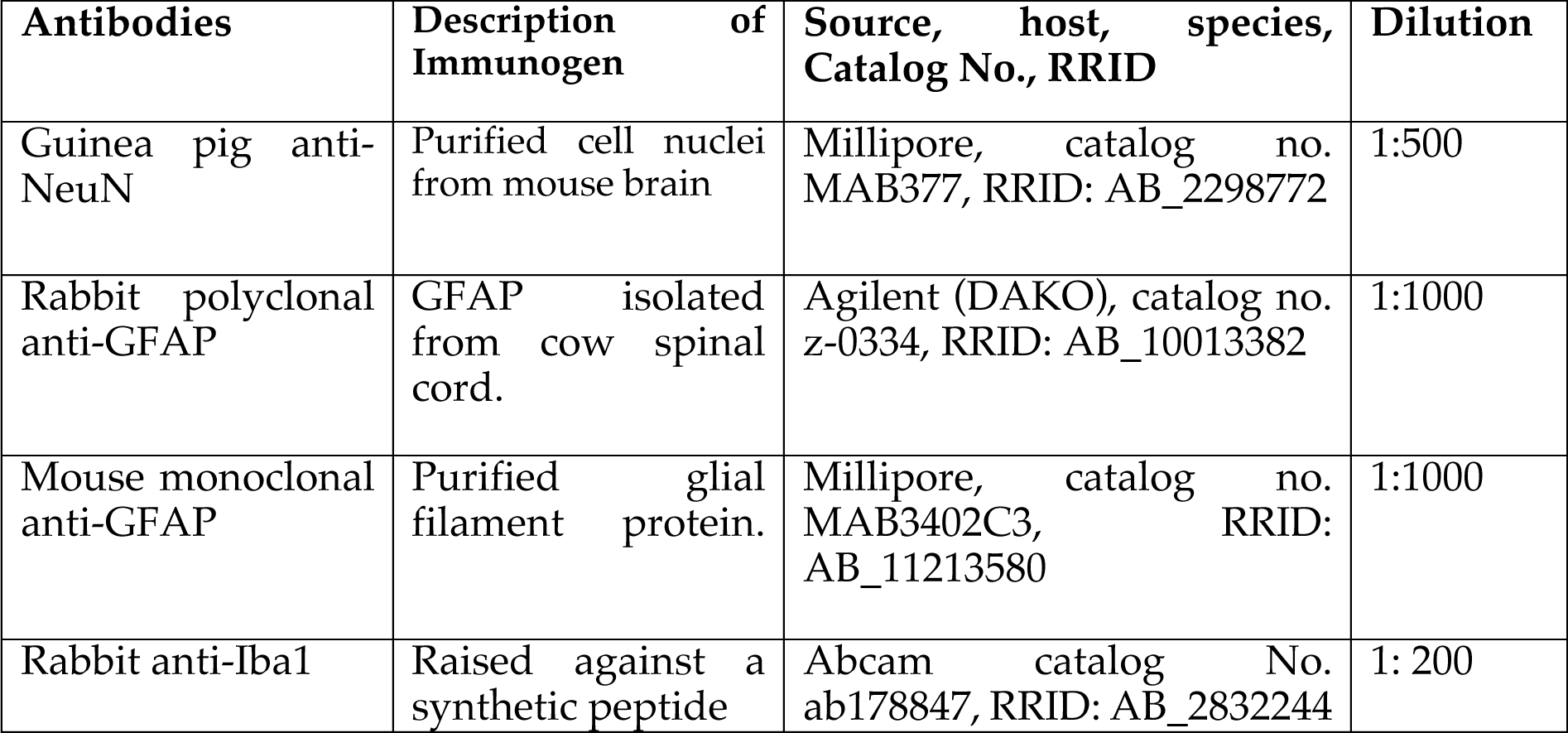
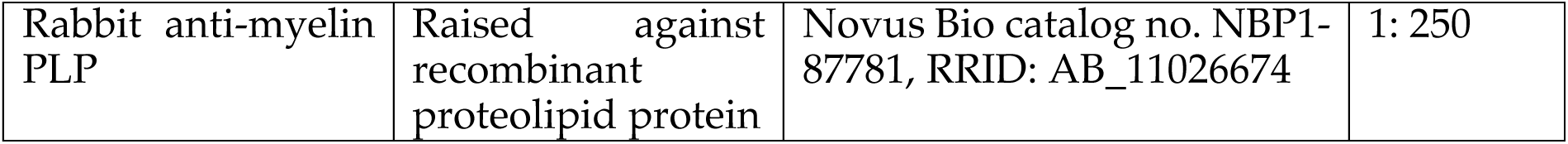
List of primary antibodies used in this study for cellular identification.

| Antibodies | Description of Immunogen | Source, host, species, Catalog No., RRID | Dilution |
| --- | --- | --- | --- |
| Guinea pig anti-NeuN | Purified cell nuclei from mouse brain | Millipore, catalog no. MAB377, RRID: AB_2298772 | 1:500 |
| Rabbit polyclonal anti-GFAP | GFAP isolated from cow spinal cord. | Agilent (DAKO), catalog no. z-0334, RRID: AB_10013382 | 1:1000 |
| Mouse monoclonal anti-GFAP | Purified glial filament protein. | Millipore, catalog no. MAB3402C3, RRID: AB_11213580 | 1:1000 |
| Rabbit anti-Iba1 | Raised against a synthetic peptide | Abcam catalog No. ab178847, RRID: AB_2832244 | 1: 200 |
| Rabbit anti-myelin PLP | Raised against recombinant proteolipid protein | Novus Bio catalog no. NBP1-87781, RRID: AB_11026674 | 1: 250 |

**Table 2.** List of secondary antibodies used in immunostaining procedures.

| Secondary antibody | Visualization for | Source, Catalog No., RRID | Dilution |
| --- | --- | --- | --- |
| Donkey Anti-Rabbit IgG (H+L) | Iba1, PLP, GFAP | Jackson ImmunoResearch, catalog no. 711005152, AB_2340585 | 1:250 |
| Donkey Anti-Guinea Pig IgG(H+L) | NeuN | Jackson ImmunoResearch, catalog no. 706005148, AB_2340443 | 1:250 |

### 2.3 Antibody characterization

The anti-NeuN antibody (Millipore, Cat# MAB377, RRID: AB_2298772) was cultivated in mice (clone A60) against the neuron-specific protein NeuN, which is expressed in most neurons in the adult mouse brain and not in glia cells (Mullen et al., 1992). This antibody’s specificity for neurons has been validated using immunohistochemistry and immunoblast analysis, showing that NeuN antibody binds to the nuclear fraction of brain tissue (Mullen et al., 1992).

The Microtubule Associated Protein 2 (MAP2) (Thermo Fisher Scientific cat. #PA1-10005, RRID: AB_1076848) was raised against recombinant constructs of the entire human projection domain, and so recognizes only the high molecular MAP2 forms, MAP2A and MAP2B. This chicken polyclonal antibody has been used to label the dendrites and perikaryal of neurons (Qiu et al., 2020; de Rus Jacquet et al., 2021).

Different anti-GFAP antibodies were used to validate the specificity of GFAP staining for astrocytes in mice (see Figure 2a-c and Table 1). These GFAP antibodies have been effectively used to immunostain astrocytes in rats, non-human primates, and humans (Forny-Germano et al., 2014; Sheikhbahaei et al., 2018a; Dominy et al., 2019; Turk and SheikhBahaei, 2022) The rabbit polyclonal anti-GFAP antibody (Agilent (DAKO), catalog #z-0334, RRID: AB_10013382) was isolated from cow spinal cord and cross-reacts with the intra-cytoplasmic filamentous protein of an epitope of the astrocytic cytoskeleton in mouse, rat, and human [manufacturer’s technical information; also see (Eng et al., 2000)]. This antibody is specific to astrocytes of the central nervous system and shows a double band (at 245–395 kDa) on Western blot (Key et al., 1993).

**Figure 2.**
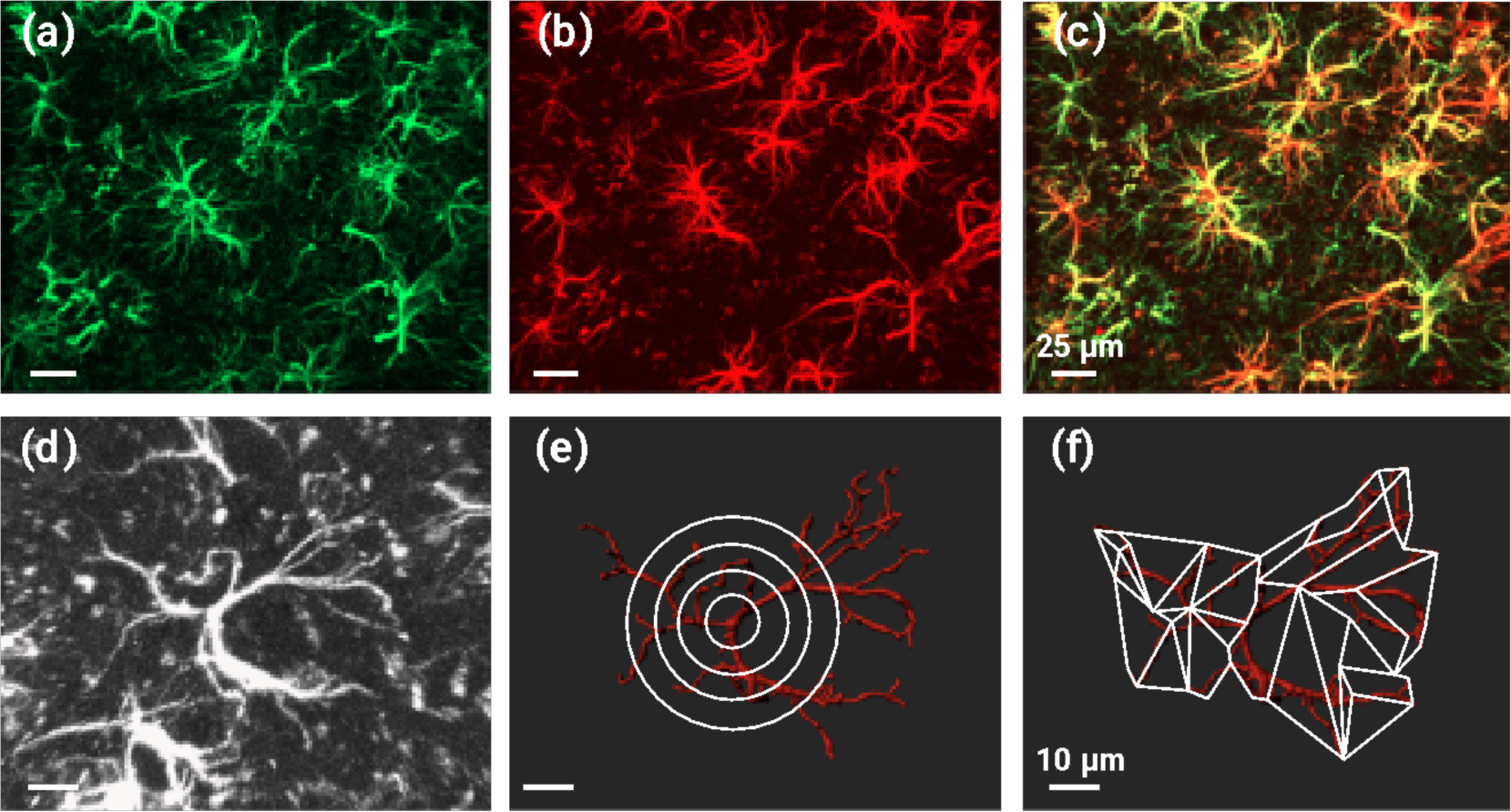
Validation of GFAP antibody and analytical methods for astrocyte morphometry. (**a – c**) Astrocytes stained with mouse anti-GFAP monoclonal antibody (green; **a**) and rabbit anti-GFAP polyclonal antibody (red; **b**). (**c**) Overlay of both staining methods highlighting colocalization of GFAP antibodies. (**d – f**) Confocal image of an astrocyte immunostained with GFAP antibody (**d**, white) and three-dimensional reconstruction of the astrocyte (**e & f**, red) using Imaris software. Sholl analysis and convex hull analysis (white lines, see Methods) are illustrated in **e** and **f**, respectively.

The mouse monoclonal GFAP antibody (MilliporeSigma, catalog# MAB3402C3, RRID: AB_11213580) was cultured against purified glial filament protein (Debus et al., 1983) and reacted with human, porcine, chicken, and rat GFAP (manufacturer’s technical information). These two anti-GFAP antibodies were used to label astrocytes in rats and non-human primates (Sheikhbahaei et al., 2018a; Turk and SheikhBahaei, 2022) and similarly found to reveal a similar pattern of labeling in GFAP-positive astrocytes in mice (Figure 2a-c).

The rabbit monoclonal Iba1 antibody (Abcam Cat #ab178846, RRID: AB_2636859) was raised against a synthetic peptide corresponding to Iba1 and reacted with Iba1 from mice, rats, and humans. The antiserum stains a single band of 15 kDa molecular weight on Western blot (manufacturer’s technical information). This rabbit monoclonal antibody has been used to label microglia (Daskoulidou et al. 2023; Mildner et al. 2017; Pajarillo et al. 2023)

The rabbit polyclonal Myelin PLP antibody (Novus Bio, Cat #NBP1-87781, RRID: AB_11026674) was raised against recombinant proteolipid protein and reacted with PLP from mouse, rat, and human (manufacturer’s technical information). According to the manufacturer, this rabbit polyclonal antibody functions as an oligodendrocyte marker (manufacturer’s technical information)

### 2.5 Three-dimensional reconstruction of astrocytes and microglia cells

Confocal z-stack images were imported into Imaris 9.7.0 (Oxford Instruments, RRID: SCR_007370), where reconstructions of individual astrocytes or microglia cells were completed with the software tracing tools. Images were optimally obtained to cover regions of interest and minimize potential overlap from other regions. Astrocytes or microglia cells with fully intact GFAP-or Iba1-immunostained processes, respectively, were chosen for reconstruction, and the cellular processes were traced throughout the entire thickness of the sections. GFAP-positive astrocytes and Iba-1-positive microglia from regions involved in vocal production [primary and secondary motor cortices (M1/M2), anterior cingulate cortex (ACC), central amygdala (CeM), and periaqueductal gray (PAG)] and control regions [secondary somatosensory cortex (S2), piriform cortex (Pir), hippocampus CA1 subregion (CA1)] were selected for reconstruction analysis (five to eight images per region across the rostral-caudal; two to four astrocytes per image) (Figure 2d-f). For cortical regions, we focused on layer V astrocytes. Astrocytic and microglia processes were traced by one investigator and verified by a second investigator. Structural quantifications of the reconstructed cells were analyzed in Imaris.

### 2.6 Morphometric analysis of astrocytes and microglia

Imaris was used to process and retrieve morphometric data from fully traced and reconstructed astrocytes and microglia cells (Turk and SheikhBahaei, 2022). The 3D filament data tool, initially developed to analyze reconstructed neurons, was utilized to carry out morphometric analysis of astrocytes and microglia cells in the regions of interest specifically. Extracted features included Sholl analysis (Sholl, 1953) to determine the unique cellular process found in each region. The complexity of astrocytic and microglial preprocesses increases with radial distance from the soma; hence, Sholl analysis quantifies process branches, branch points, number of terminal points, as well as process length in astrocytes and microglia cells. This analysis utilized shell volumes between concentric spheres, each 1 μm apart, radiating out from the center of the soma (see Figure 2e). Notably, the Sholl analysis locates the number of intersections between processes and spheres at a given radius.

In astrocytes, the processes branch from primary to secondary, tertiary, branchlets, and even fine leaflets, which may be unrecognizable with GFAP staining. Because of the complexity of astrocyte morphology, the 3D convex hull analysis was used to assess the volume occupied by the astrocytic process as described before (Sheikhbahaei et al., 2018a; Turk and SheikhBahaei, 2022). In convex hull analysis, the volume and area of astrocytes were *estimated* by enveloping the cell surface area and volume, creating a polygon that joins terminal points of the processes (see Figure 2f).

We then used the complexity index (CI) to normalize the comparison of the overall cellular complexity in distinct regions. This CI has been used to evaluate the complexity of neurons (Diamantaki et al., 2016; Guillamon-Vivancos et al., 2019; Reagan et al., 2021) and glia cells (Sheikhbahaei et al., 2018a; Turk and SheikhBahaei, 2022). CI was computed by using the following formula: (Σ terminal orders + number of terminals) × (total process length/number of primary branches), where number of terminal orders for each terminal point is calculated as the number of branches that appear proceeding backward from the defined terminal to the cell soma (Sheikhbahaei et al., 2018a)

### 2.7 Percentage of immunostained area for neurons, oligodendrocytes, and white matter astrocytes

Cellular processes of neurons and oligodendrocytes were stained with Map2 and PLP, respectively (Table 1). Due to significant overlaps, the processes of immunostained individual neurons or oligodendrocytes were difficult to identify; therefore, to quantify differences in morphological characteristics of these cell types, the percent staining area in each region of interest was measured between *Gnptab*-mutant and control animals. Image analysis was performed with ImageJ-Fiji software (RRID: SCR_002285) (Schindelin et al., 2012), in which a minimum intensity threshold was set to characterize PLP- or Map2 immunostaining.

Similarly, astrocytes in the corpus callosum branch extensively from their primary processes, resulting in significant overlap between processes. Because of this complex overlap, we were not able to reconstruct their and instead used ImageJ-Fiji to calculate the percent area of GFAP-positive stain in this region structure (Meyer et al., 2018; Molina-Gonzalez et al., 2023). A minimum intensity threshold was used to define GFAP-positive processes.

### 2.6 Statistical analysis

The analyzed data were exported from Imaris to Prism 9.0 software (Graphpad Software Inc., RRID: SCR_002798) for statistical analysis, where the data were reported as averages ± standard error of the mean (SEM). We used non-parametric Mann–Whitney U rank test or the Kruskal-Wallis one-way ANOVA by ranks followed by Dunn’s post hoc as appropriate for statistical analysis.

## 3 RESULTS

### 3.1 Analysis of neuronal processes

We first analyzed the distribution of neuronal processes in brain regions involved in vocal production and control regions (Figure 3) in *Gnptab*-mutant mice and control littermates. We used NeuN and MAP2 immunostaining (Figure 3a&b) to identify neurons’ cell bodies and processes. Since MAP2-stained processes of neurons were densely cross-covered, we could not precisely trace the processes of individual neurons. As a result, we did not further investigate the morphology of neurons but rather analyzed the MAP2-immunostained area. We found no differences in the distribution of neuronal processes in primary and secondary motor cortices (M1/M2), anterior cingulate cortex (ACC), piriform cortex (Pir), secondary somatosensory cortex (S2), periaqueductal gray (PAG) central amygdala (CeM), and hippocampal CA1 regions between *Gnptab*-mutant and control mice (Figure 3c).

**Figure 3.**
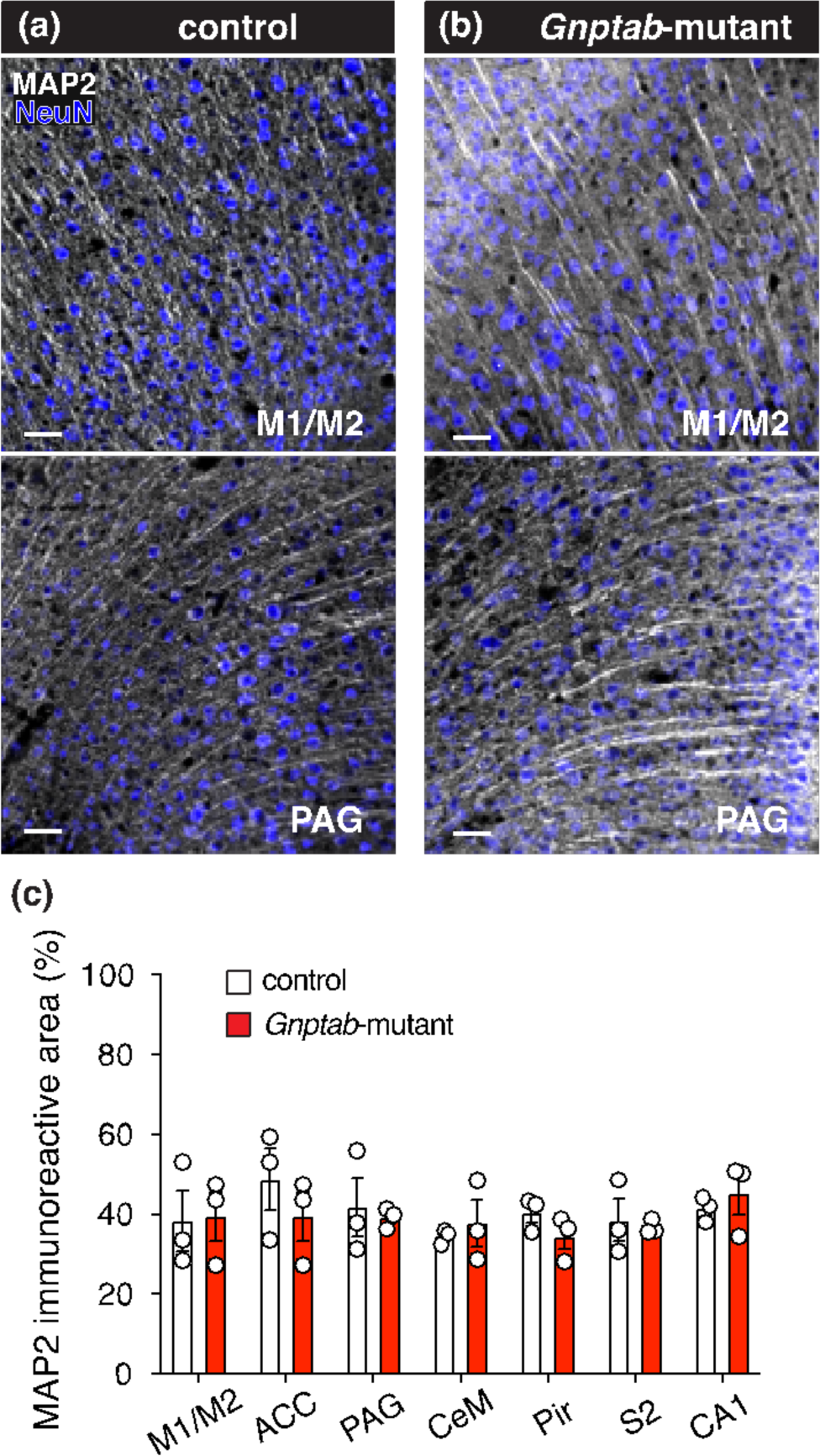
Immuno-stained MAP2-positive neurons in the adult *Gnptab*-mutant and control mice. (**a & b**) Confocal imagery of neurons immunostained with MAP2 (*white*) and NeuN (*blue*) in primary and secondary motor cortices (M1/M2; *top*) and periaqueductal gray (PAG; *bottom*) in control (**a**) and *Gnptab*-mutant (**b**) mice. Scale bars: 25 µm. (**c**) Summary data illustrating the distribution of MAP2-positive neurons (as % of surface area) from control (n = 3) and *Gnptab*-mutant (n = 3) mice. ACC, anterior cingulate cortex; CeM, central nucleus of the amygdala; Pir, piriform cortex; S2, secondary somatosensory cortex; CA1, hippocampus CA1.

**Figure 4.**
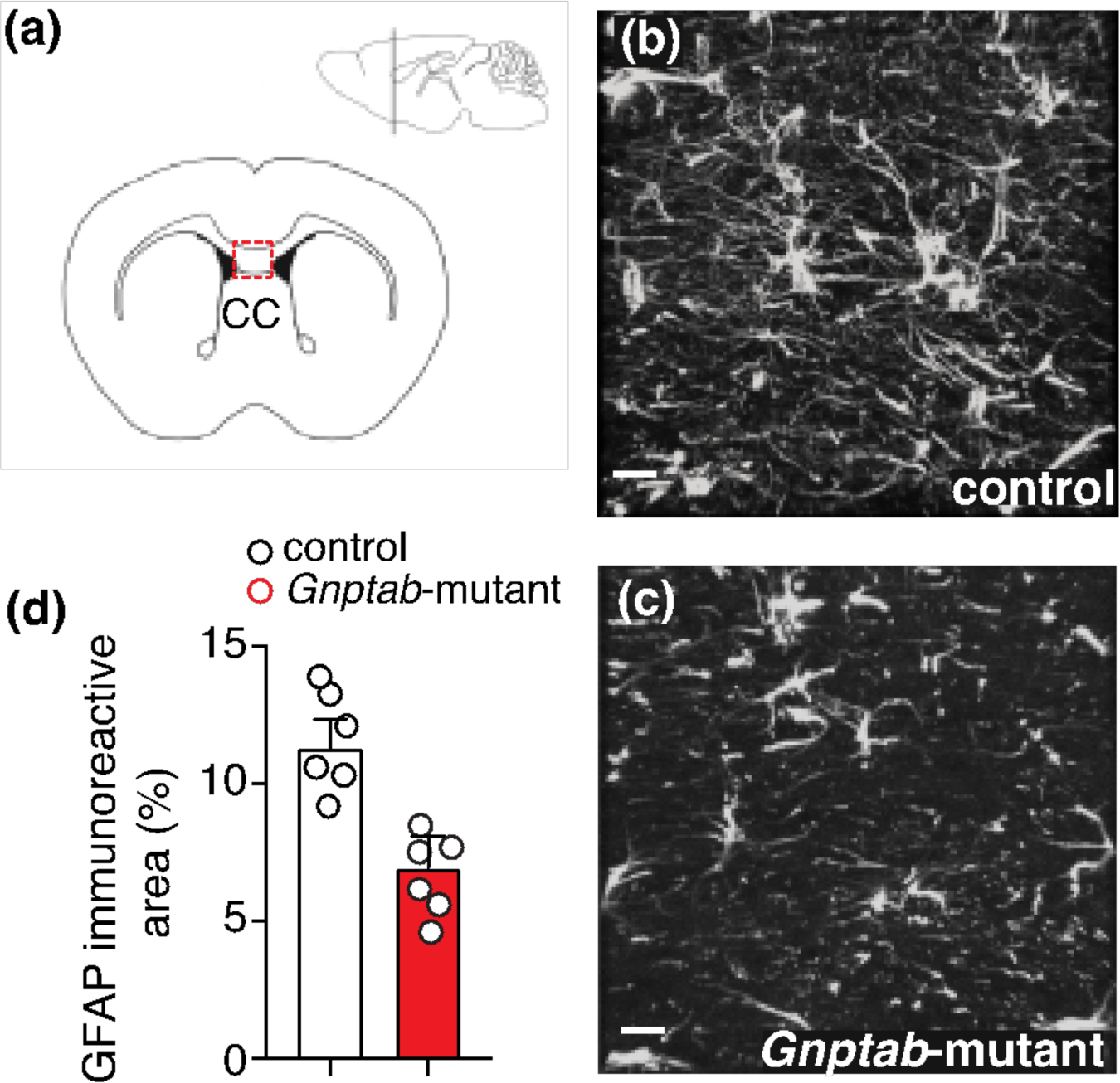
Effect of mutation in *Gnptab* on GFAP-positive astrocytes in the corpus callosum. **(a)** Sagittal and coronal illustration of the regions of corpus callosum (CC) in adult mice. The red dashed box depicts the genu of CC, our region of interest**. (b & c)** Confocal images of GFAP-positive astrocytes (white) in the genu of CC in control (**b**) and *Gnptab*-mutant (**c**) mice. Scale bars: 20 µm. **(d)** Group data illustrating the distribution of GFAP-positive astrocytes (as % of surface area) from control (n = 6) and *Gnptab*-mutant (n = 6) mice.

### 3.2 Analysis of corpus callosum astrocytes

Previous reports indicated that astrocytes in the corpus callosum were impacted in the *Gnptab*-mutant mice (Han et al., 2019). Cell bodies of these relatively sparse white-matter astrocytes were found to be inside the corpus callosum and have numerous long processes coursing in the mediolateral plane, creating a prominent network of astrocytic fibers (Figure 3). The densely intermingled GFAP-stained processes of callosal astrocytes prevented precise tracing of individual astrocyte processes. As a result, we did not further assess the morphology of callosal astrocytes but rather analyzed the GFAP-immunostained area at the genu of the corpus callosum (Figure a-c). Consistent with the previous report (Han et al. 2019), we found that compared to the control mice, the GFAP-stained area was decreased (by 39%; p = 0.002, Mann-Whitney U test; n = 6) in the corpus callosum of *Gnptab*-mutant mice (Figure d).

### 3.3 Morphometric analysis of astrocytes

Morphometric evaluation was carried out on reconstructed GFAP-positive astrocytes in cortical regions to delineate specific morphological properties of these star-shaped brain cells in *Gnptab*-mutant and control littermate animals (Figure 5). Sholl and convex hull analyses were used to quantify branch points, number of process terminal points, length of processes (Figures 2&5), number of process intersections, as well as convex hull volume and surface area of the reconstructed astrocytes from M1/M2, ACC, Pir, and S2. We found that astrocytes residing in cortical regions associated with vocal production (i.e., M1/M2 and ACC) were smaller and less complex when compared to those in the control mice (Figure 6; Table 3). Similarly, when comparing the morphometric characteristics of midbrain astrocytes brain regions that are involved in vocal production (i.e., PAG and central amygdala CeM), we found that the *Gnptab*-mutant astrocytes were again smaller and less complex when compared to those in the control mice (Figure 6; Table 3). We did not find differences in the cellular morphology of astrocytes in the hippocampal CA1 region (Figure 6; Table 3). To normalize and compare the complexity of cell processes among astrocytes from various regions, we employed the Complexity Index (CI), which has been used for the study of astrocyte morphology (Sheikhbahaei et al., 2018a; Turk and SheikhBahaei, 2022). In the *Gnptab*-mutant mice, astrocytes residing within the M1/M2, ACC, CeM, and PAG showed lower CI compared to the ones in control mice (Figure 6f; Table 3), suggesting that astrocytes in the *Gnptab*-mutant mice are less complex compared to the control animals. Together, these data indicate that this specific mutation in the *Gnptab* gene directly affects astrocytes in the brain regions associated with motor behaviors of vocalization.

**Figure 5.**
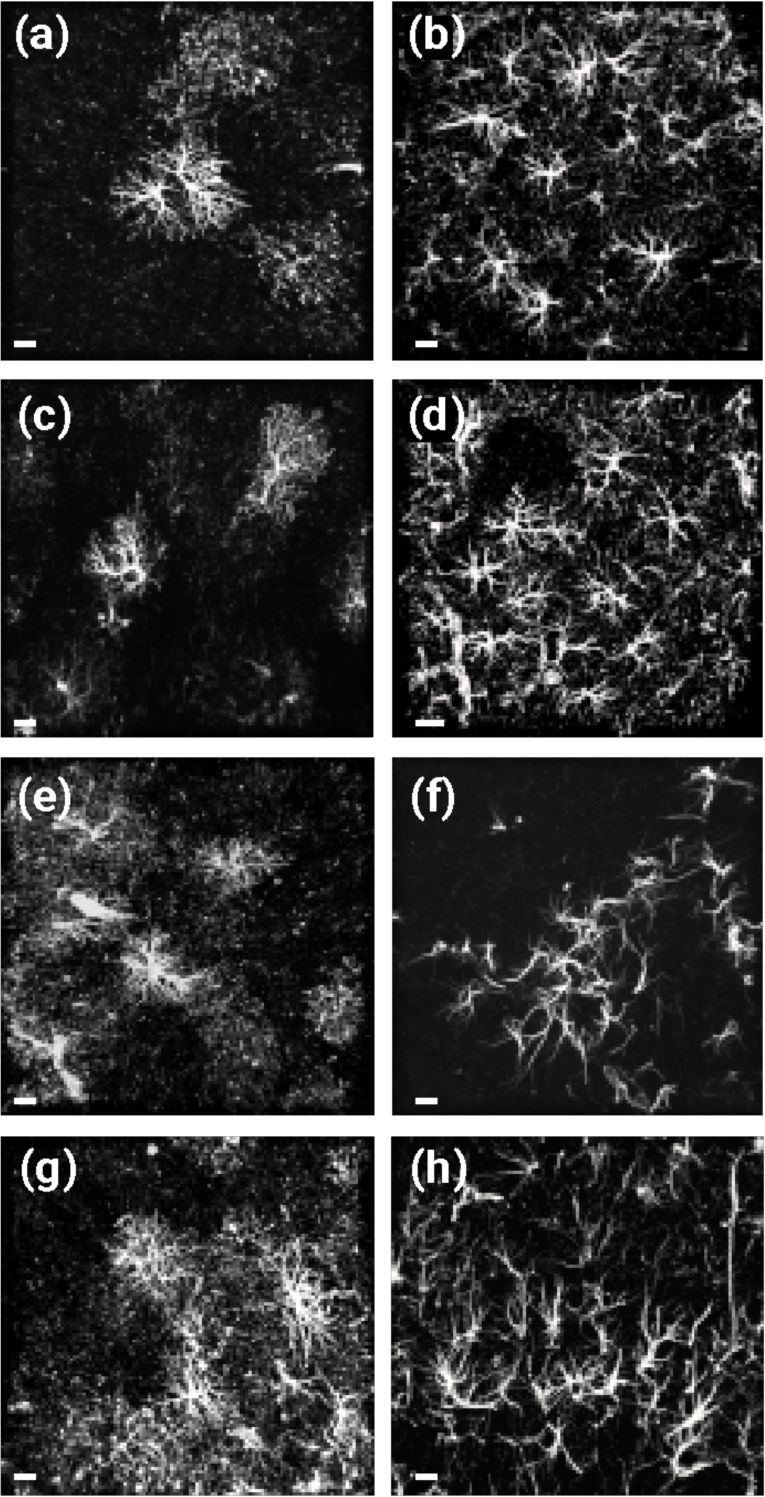
Immunostained GFAP-positive astrocytes in the adult mouse brain. Confocal images of astrocytes (white) immunofluorescently labeled with GFAP antibody in regions of interest (see Figure 1), including the primary motor cortex (M1, **a**), the secondary motor cortex (M2, **b**), anterior cingulate cortex (ACC, **c**), the secondary somatosensory cortex (S2, **d**), central nucleus of the amygdala (CeM, **e**), periaqueductal gray (PAG, **f**), piriform cortex (Pir, **g**), and hippocampus CA1 **(h)**. Scale bars: 10 µm.

**Figure 6.**
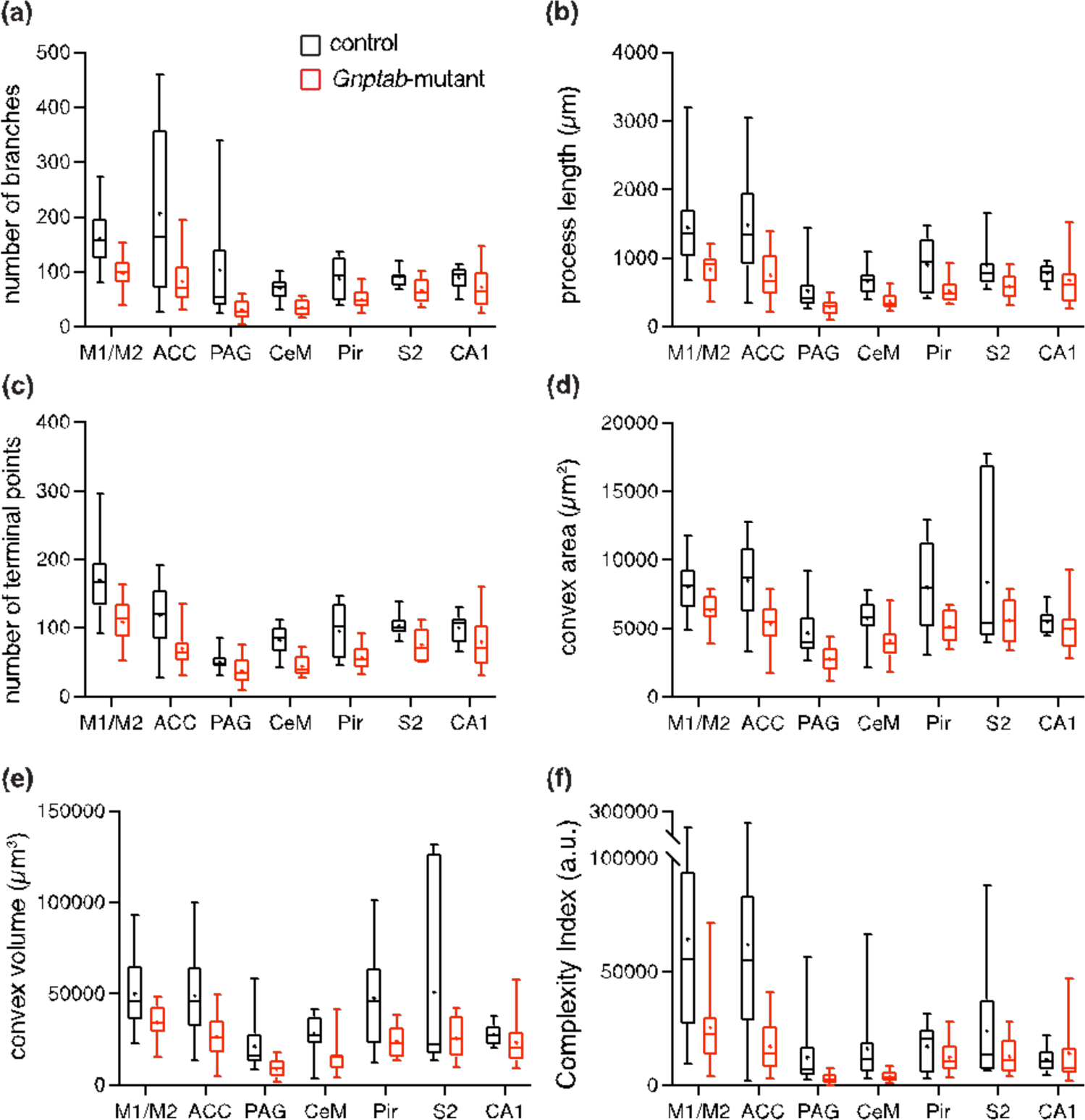
Morphometric features of astrocytes in *Gnptab*-mutant mice. Summary data illustrating morphometric analysis of data obtained using sholl analysis: (**a**) number of branches, (**b**) process length, (**c**) number of terminal points, (d) convex hull surface area, (**e**) convex hull volume. (**f**) Comparison of normalized data from each region of interest using complexity index (see Methods). Overall, astrocytes residing within vocal motor circuits were less complex in *Gnptap*-mutant mice than in control littermates (see Table 3 for more details). M1/M2, primary and secondary motor cortices; ACC, anterior cingulate cortex; PAG, periaqueductal gray; CeM, central nucleus of the amygdala; Pir, piriform cortex; S2, secondary somatosensory cortex; CA1, hippocampus CA1. *n* = 10 – 20 astrocytes per region.

### 3.4 Morphometric analysis of microglia

Sholl analysis was used to investigate the morphometric characteristics of reconstructed microglia from the cortical and midbrain regions in the *Gnptab*-mutant and control mice (Figure 7). We then compared the average number of branch points, terminal points, and process length from the reconstructed microglia cells in each brain region between the *Gnptab*-mutant and control mice (Figure 8; Table 4). Our data suggest robust differences in all morphometric parameters from Iba1-reconstructed microglia in only M1/M2 and ACC (Figure 8; Table 4). Microglia cells in these regions had fewer branch and terminal points and shorter processes (Figure 8; Table 4). Accordingly, the CI of reconstructed M1/M2 and ACC microglia cells from *Gnptab*-mutant mice was lower than that of the control (Figure 8f; Table 4). These data suggest that the Gnptab gene mutation primarily affects microglia cells’ morphological characteristics in cortical regions associated with vocal production but not in other vocal production regions (i.e., CeM and PAG) or control regions (i.e., S2 and Pir).

**Figure 7.**
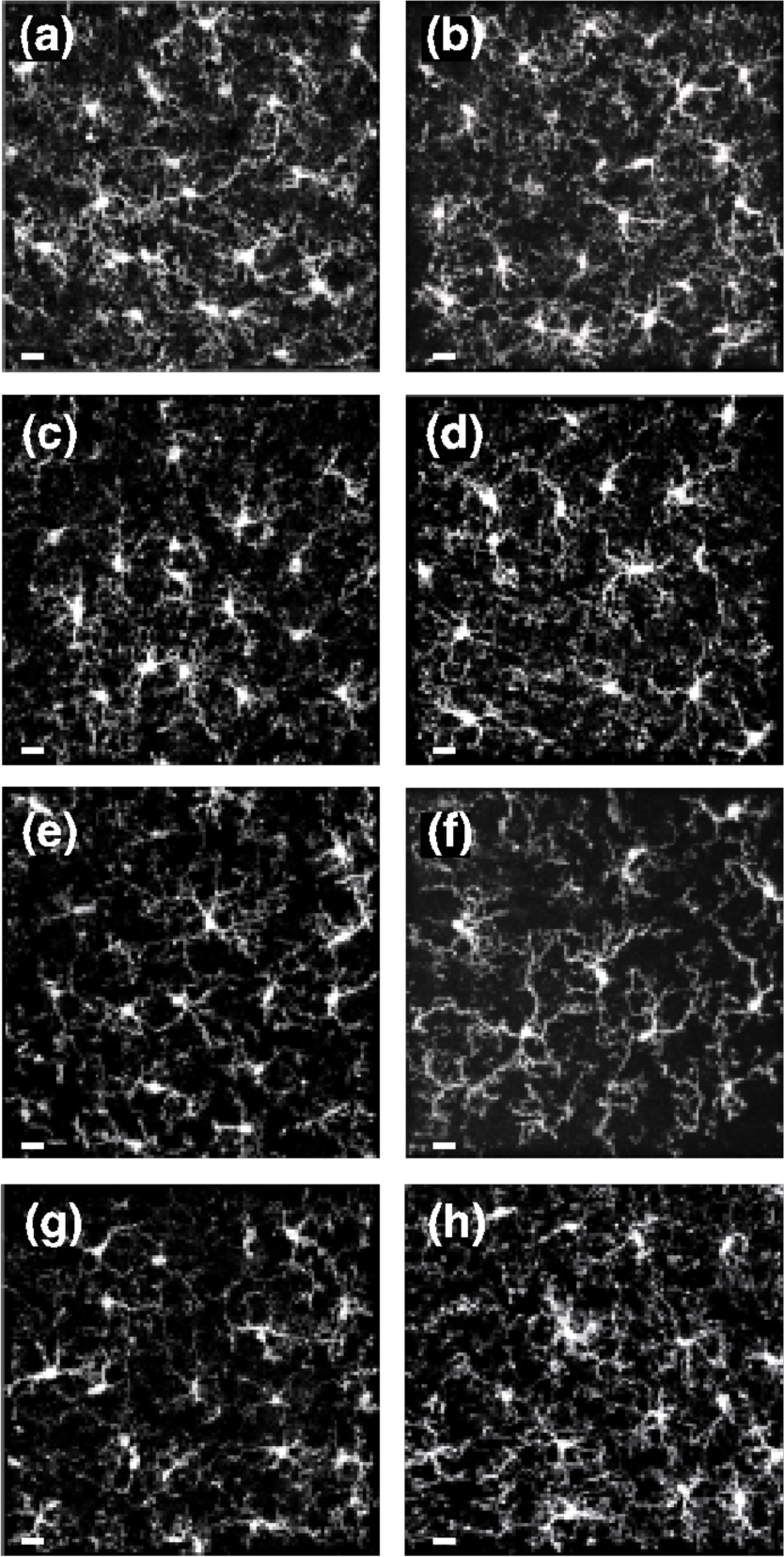
Immunostained Iba1-positive microglia cells in the adult mouse brain. Confocal images of immunolabelled Iba1-positive microglia (white) in regions of interest (see Figure 1), including primary motor cortex (M1, **a**), secondary motor cortex (M2, **b**), anterior cingulate cortex (ACC, **c**), secondary somatosensory cortex (S2, **d**), central nucleus of the amygdala (CeM, **e**), periaqueductal gray (PAG, **f**), piriform cortex (Pir, **g**), and hippocampus CA1 **(h)**. Scale bar: 10 µm.

**Figure 8.**
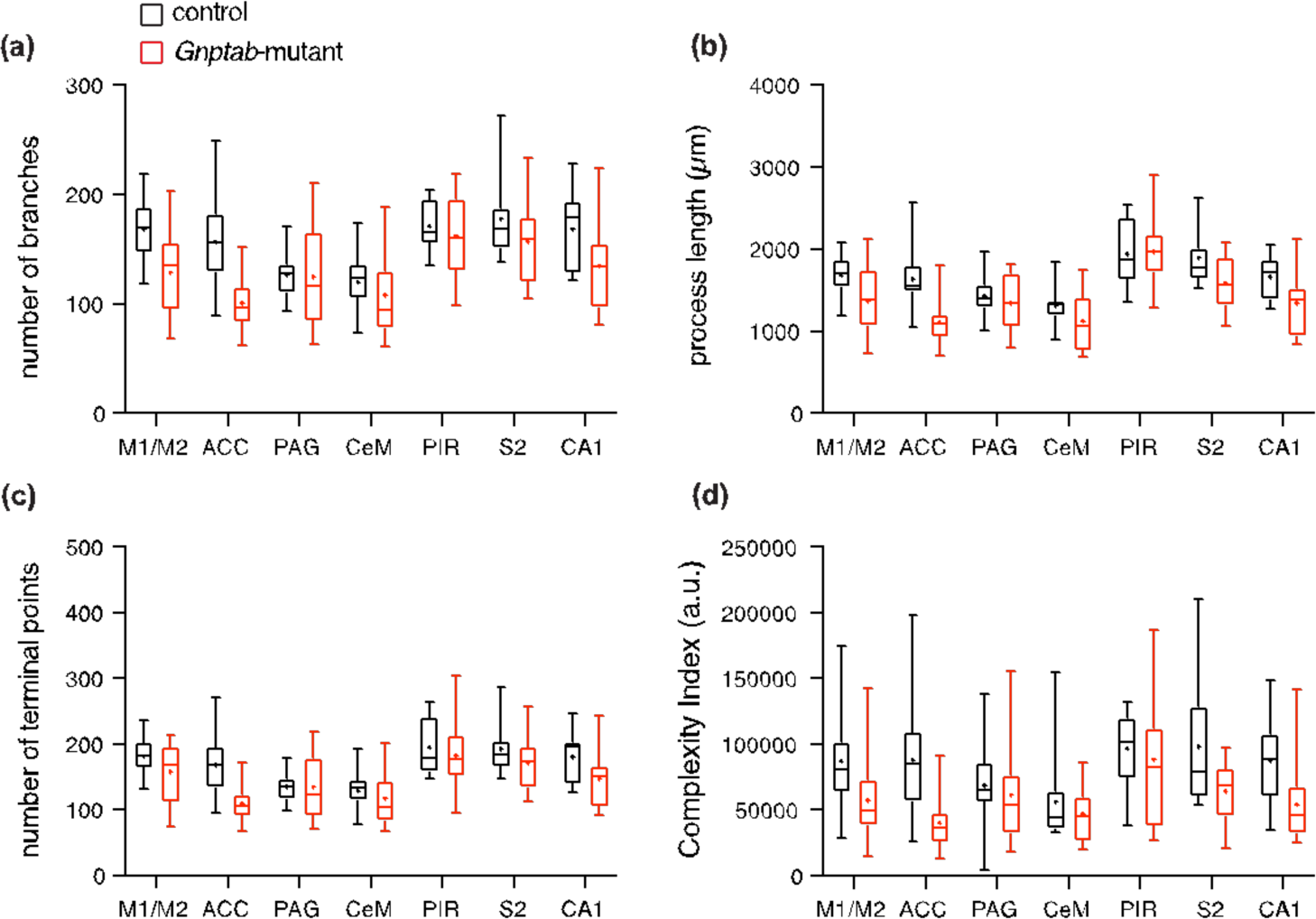
Morphometric characteristics of microglia in *Gnptab*-mutant mice. Summary data illustrating the morphometric analysis of data obtained using Sholl analysis: (**a**) number of branches, (**b**) process length, and (**c**) number of terminal points. (**f**) To normalize and compare data from each region, we used the complexity index (see Methods). Overall, microglia residing within cortical vocal motor circuits were less complex in *Gnptap*-mutant mice than in control littermates. See Table 4 for more details. M1/M2, primary and secondary motor cortices; ACC, anterior cingulate cortex; PAG, periaqueductal gray; CeM, central nucleus of the amygdala; Pir, piriform cortex; S2, secondary somatosensory cortex; CA1, hippocampus CA1. *n* = 10 – 20 microglia per region.

### 3.5 Analysis of oligodendrocytes

We used an anti-proteolipid protein (PLP) antibody to immunostain oligodendrocyte cells (Timsit et al., 1995) (Figure 9a&b). Like corpus callosum astrocytes, we could not reconstruct individual cellular processes of oligodendrocytes because of the long and intermingled nature of processes. Similarly, we compared the percent surface area of PLP staining in regions of interest between *Gnptab*-mutant and control mice. We did not identify any differences of PLP-stained areas between two experimental groups (Figure 9c; Table 5).

**Figure 9.**
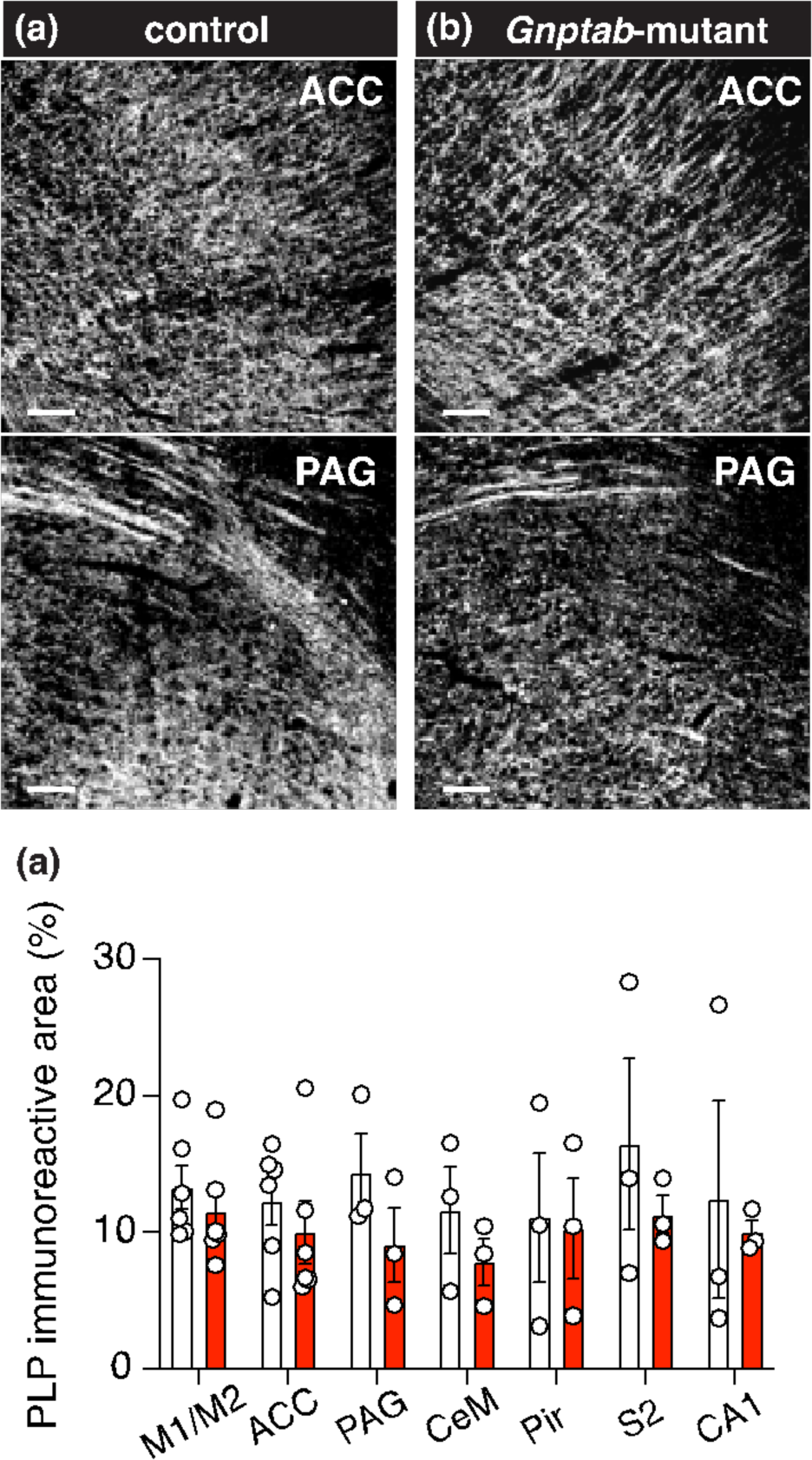
Immuno-stained PLP-positive oligodendrocytes in the adult *Gnptab*-mutant and control mice. (**a & b**) Confocal imagery of neurons immunostained with PLP (*white*) in the anterior cingulate cortex (ACC; *top*) and periaqueductal gray (PAG; *bottom*) in control (**a**) and *Gnptab*-mutant, (**b**) mice. Scale bars: 25 µm. (**c**) Group data illustrating the distribution of PLP-positive oligodendrocytes (as % of surface area) from control and *Gnptab*-mutant mice (n = 3 – 6 mice per group). M1/M2, primary and secondary motor cortices; CeM, central nucleus of the amygdala; Pir, piriform cortex; S2, secondary sensory cortex; CA1, hippocampus CA1.

## 4 DISCUSSIONS

Although the precise neural circuits governing vocal production have not yet been elucidated, it is widely acknowledged that specific cortical and midbrain regions [such as primary motor cortex (M1), supplementary motor cortex (pre-SMA), anterior cingulate cortex (ACC), central nucleus of amygdala (CeM), and periaqueductal gray (PAG)] partake in orchestrating this intricate behavior (Jürgens, 2002; Jarvis, 2019). While innate vocalization might engage midbrain regions, including CeM and PAG, voluntary vocalization, deemed a cognitive function, entails a spectrum of cortical regions, encompassing frontal regions including M1, pre-SMA, and ACC (Petrides, 2005; Simões et al., 2010; Burman et al., 2014; Miller et al., 2015; Theodoni et al., 2020).

In this study, we probed the structural attributes of neurons and glia cells within the vocal production circuits of adult mice in a genetically modified mouse model with a *Gnptab* mutation (Barnes and Holy, 2016), which is associated with human stuttering (Kang et al., 2010). Our data suggest that the cytoarchitecture of astrocytes and microglia cells in the vocal production circuits was affected by the *Gnptab* mutation in mice.

### 4.1 Developmental stuttering and use of animal models

Developmental stuttering (stuttering) is a neurodevelopmental disorder characterized by disruptions in speech timing and fluency, often manifesting as repetition, prolongation, or pausing of words (American Psychiatric Association, 2013; Bloodstein et al., 2021).

Stuttering may be associated with motor tics, cognitive avoidance, social anxiety, and notable comorbidity with attention-deficit/hyperactivity disorder (ADHD) (Kraaimaat et al., 2002; Donaher and Richels, 2012; Maguire et al., 2020). Emerging predominantly in early childhood, it affects around 5-8% of children, with over 1% continuing to experience symptoms into adulthood (Yairi and Ambrose, 2013). The etiologies of stuttering encompass genetic predispositions, abnormal basal ganglia development, and alterations in white matter tracts (Alm, 2004; Lu et al., 2010; Craig-McQuaide et al., 2014; Chang and Guenther, 2019; Maguire et al., 2021; Mollaei et al., 2021; SheikhBahaei et al., 2022). Neuroimaging studies in humans have shown lower activity within the left hemispheric speech domains and reduced brain activity in the left frontal precentral cortex in individuals with stuttering (Pool et al., 1991; Watkins et al., 2008; Chang et al., 2011; Beal et al., 2013).

Although mutations in several genes are linked to developmental stuttering (Kang et al., 2010; Shaw et al., 2021; Polikowsky et al., 2022; Below et al., 2023), few of them have been studied with animal models nor been shown to have causative effects (Kang et al., 2010; Barnes et al., 2016). Animal models have been used to study mechanistic questions in other neurodevelopmental. As such, a mouse model remains of great interest for studying developmental stuttering at the cellular and circuit levels.

### 4.2 Neuron pathology

The etiology of stuttering is generally unknown, but it is established that stuttering is a neurodevelopmental motor control disorder (Garnett et al., 2018; Kim et al., 2020; Höbler et al., 2022). Pathologies in neuronal cells/circuits have been identified in other neurodevelopmental disorders(Sahin and Sur, 2015; Cravedi et al., 2017; Del Pino et al., 2018; Krol et al., 2018), however, regardless of stuttering severity, people who stutter may speak fluently from time to time. This observation suggests that at least in a group of individuals who stutter, the fundamental neuronal circuits generating speech are intact, and the symptom may instead arise in the coordination of activities of brain circuits critical for speech production and/or other cells coordinating activities of neurons (i.e., glia cells). Consistent with this hypothesis, we did not identify differences in neuronal processes in the vocal production circuits between the *Gnptab*-mutant and control littermates (Figure 3). However, this morphological analysis does not assess the activities of neurons and more experiments are needed to assess neuronal activities in the brain circuits involved in vocal production. On the other hand, it has been proposed that, at least in a group of people who stutter, the deficit in developmental stuttering might arise in glial cells (Han et al., 2019; Maguire et al., 2021; Turk et al., 2021; Frigerio Domingues et al., 2022); as such, we then systematically studied morphometric characteristics of glial cells.

### 4.3 Glial pathology: astrocytes

Previous studies in the mouse model employed here suggested that the corpus callosum astrocytes were affected by the *Gnptab* mutation (Han et al., 2019); however, detailed analysis of the cytoarchitecture of astrocytes (and other glia cells) in the vocal production circuits were lacking.

Astrocytes, specialized glial cells in the Central Nervous System (CNS), play a key role in maintaining brain homeostasis (Halassa et al., 2009; Allaman et al., 2011; Marina et al., 2018). Historically relegated to a supporting role, recent studies have highlighted astrocytes’ diverse functions, underscoring their importance in synaptic activity modulation, neuronal functional maintenance, including regulation of extracellular pH and K^+^ levels, and neurotransmitter metabolism (Iadecola and Nedergaard, 2007; Gordon et al., 2008). Additionally, they are crucial for neuronal plasticity, synapse formation, and activity regulation, further contributing to the modulation of brain cells and circuits through gliotransmitters (Araque et al., 2014; Sheikhbahaei et al., 2018b; Bishop and SheikhBahaei, 2023; SheikhBahaei et al., 2023a)

It has been shown that astrocytes have modulatory roles in other motor circuits (Sheikhbahaei et al., 2018b; Turk et al., 2022) affecting complex behaviors. Therefore, we hypothesize that astrocytes may also modulate vocal motor circuits, and a deficit in these astrocytes might lead to stuttering. Therefore, we explored the morphometric properties of astrocytes from both cortical and midbrain regions involved in vocal production (as well as control regions) in *Gnptab*-mutant mice and control littermates. We employed antibodies against glial fibrillary acidic protein (GFAP) protein to delineate cellular structures of astroglia. GFAP, a primary constituent of glial filaments in mature astrocytes of the CNS, is a hallmark of mature astrocytic cell differentiation (Dahl et al., 1981; Bovolenta et al., 1984), and GFAP immunostaining has been used to study morphometric characteristics of rodents’ and non-human primates’ astrocytes (Saur et al., 2014; Sheikhbahaei et al., 2018a; Turk and SheikhBahaei, 2022) After validation of GFAP antibodies in mice, we compared the morphometric characteristics of astrocytes in the corpus callosum, brain regions that are hypothesized to be involved in mice vocal production (i.e., M1/M2, ACC, CeM, and PAG), and a few control regions (CA1, S2, Pir) in *Gnptab*-mutant mice and control littermates.

Consistent with previous reports (Sofroniew and Vinters, 2010; Khakh and Sofroniew, 2015), the number of GFAP-stained astrocytes in the cortical regions was low in both *Gnptab*-mutant and control mice. Therefore, in M1, M2, ACC, S2, and Pir cortices, we reconstructed astrocytes across the rostral-caudal axis to increase the number of imaged and reconstructed astrocytes. In this context, we did not characterize the regional specificity of cellular properties of cortical astrocytes along the imaging axis. Future experiments will be needed to define the cytoarchitecture of astrocytes in M1, M2, and ACC in different rostral-caudal regions, as they may have different projections.

### 4.4 Glia pathology: Microglia

Microglia, the resident phagocytes of the CNS, are essential for maintaining homeostasis during development and in neurodevelopmental disorders (Kettenmann et al., 2011; Paolicelli et al., 2011). They transition from mediating developmental inputs to modulating neuronal activity in adulthood (Parkhurst et al., 2013). Microglia cells sense brain metabolites, influencing neuronal activity and homeostasis (Schafer et al., 2012). They are also integral to synaptogenesis and synaptic plasticity, producing complement-related proteins that facilitate the pruning of unwanted synapses during development (Nimmerjahn et al., 2005; Wake et al., 2009; Schafer et al., 2012). The multifaceted roles of microglia in both synaptic and homeostatic regulation reflect their critical role in the development and function of CNS across the lifespan (Paolicelli et al., 2011).

Our data suggest that in addition to astrocytes, microglia cells might also be involved in the pathophysiology of stuttering. While astrocytes have been demonstrated to modulate motor circuit activities (Morquette et al., 2015; Sheikhbahaei et al., 2018b), evidence suggesting that microglia possess a similar function is limited. Therefore, we posit that microglia’s role in the pathophysiology of stuttering might be different than that of astrocytes. Notably, there’s a significant surge in postnatal synapse formation in circuits involved in vocal production, such as corpus callosum, thalamocortical pathways, and descending cortical tracts (Stanfield et al., 1982; Stanfield and O’Leary, 1985; Gilmore et al., 2007), given that the cerebral cortex establishes the majority of its synaptic connections postnatally. Given the role of microglia in synaptic refinement procedures after this surge, dysfunction in *Gnptab*-mutant mice’s microglial cells could lead to compromised connectivity in the vocal production circuits.

### 4.5 Glia pathology: Oligodendrocytes

Oligodendrocytes play a pivotal role in the function of motor circuits primarily through myelination, which enables the fast propagation of action potentials along the ensheathed axons (Kuhn et al., 2019; Moore et al., 2020; Krämer-Albers and Werner, 2023). Myelination by oligodendrocytes is fundamental for the effective operation of motor circuits, facilitating and enhancing the propagation velocity of action potentials along axons, thereby ensuring timely and synchronized motor responses (Almeida and Lyons, 2017). Additionally, oligodendrocytes contribute to the structural and functional integrity of motor circuits via myelination, aiding in the precise relay of motor commands from the central nervous system to peripheral effector organs. Beyond their myelinating role, oligodendrocytes contribute to axonal trophic support and exhibit immunomodulatory capacities, which may hold significance in the broader context of motor circuit health (Bradl and Lassmann, 2010; Stadelmann et al., 2019). Collectively, the contributions of oligodendrocytes are instrumental in sustaining the functionality and robustness of motor circuits. Therefore, we decided to study the morphology of mature oligodendrocytes using PLP immunostaining. Since intermingled processes of neighboring PLP-positive oligodendrocytes make the 3D reconstruction inaccurate, we measured the percentage of surface areas occupied by PLP-positive oligodendrocytes. Although we did not identify any differences in the expression of PLP between *Gnptab*- mutant and control mice (Figure 9), it is still possible that the function of oligodendrocytes might be affected by the mutation.

### 4.6 Outlook

Developmental stuttering is a neurodevelopmental disorder hypothesized to affect motor control circuits involved in vocal production. Our data from the *Gnptab*-mutant mouse model for stuttering suggest that, at least in a subgroup of people who stutter, the cytoarchitecture of microglia and astrocytes in vocal motor circuits are affected. We hypothesize that these glial cells are involved in the pathophysiology of stuttering via different mechanisms. Since astrocytes maintain an important function in metabolic support of neuronal activities, our data further support the hypothesis that metabolic pathways are affected in stuttering (Chow et al., 2020; Alm, 2021; Turk et al., 2021). Given the close relationship between cellular function and morphology, the function of astrocytes and microglia may also be affected in the *Gnptab*-mutant mice, a hypothesis that requires further evaluation. Although many questions remain unanswered, our data strengthen the hypothesis that dysfunction of glial cells might lead to stuttering disorder (Han et al., 2019; Maguire et al., 2021; Turk et al., 2021; Frigerio Domingues et al., 2022). Future investigations into the functional implications of the mutation in the *Gnptab* gene within these glial cells may shed light on the pathophysiological underpinnings of developmental stuttering.

## Acknowledgments

We thank NINDS Light Microscopy Core for technical support and Ariana Z. Turk for comments on the previous version of the manuscript and assistance with cellular reconstructions. We also thank Ana Paz and Demi Lombardo for administrative support. This work was supported by the Intramural Research Program of the NIH, NINDS (ZIA NS009420 to SSB).

## References

1. Allaman I, Bélanger M, Magistretti PJ. 2011. Astrocyte-neuron metabolic relationships: for better and for worse. Trends Neurosci 34:76–87.

2. Almeida RG, Lyons DA. 2017. On myelinated axon plasticity and neuronal circuit formation and function. J Neurosci 37:10023–10034.

3. Alm PA. 2004. Stuttering and the basal ganglia circuits: a critical review of possible relations. J Commun Disord 37:325–369.

4. Alm PA. 2021. Stuttering: A disorder of energy supply to neurons? Front Hum Neurosci 15:662204.

5. Ambrose NG, Yairi E, Cox N. 1993. Genetic aspects of early childhood stuttering. J Speech Hear Res 36:701–706.

6. American Psychiatric Association. 2013. Diagnostic And Statistical Manual Of Mental Disorders, 5th Edition: Dsm-5. 5th ed. Washington, D.C: American Psychiatric Publishing.

7. Araque A, Carmignoto G, Haydon PG, Oliet SHR, Robitaille R, Volterra A. 2014. Gliotransmitters travel in time and space. Neuron 81:728–739.

8. Barnes TD, Holy TE. 2016. Knockout of Lysosomal Enzyme-Targeting Gene Causes Abnormalities in Mouse Pup Isolation Calls. Front Behav Neurosci 10:237.

9. Barnes TD, Wozniak DF, Gutierrez J, Han T-U, Drayna D, Holy TE. 2016. A Mutation Associated with Stuttering Alters Mouse Pup Ultrasonic Vocalizations. Curr Biol.

10. Beal DS, Gracco VL, Brettschneider J, Kroll RM, De Nil LF. 2013. A voxel-based morphometry (VBM) analysis of regional grey and white matter volume abnormalities within the speech production network of children who stutter. Cortex 49:2151–2161.

11. Below J, Polikowsky H, Scartozzi A, Shaw D, Pruett D, Chen H-H, Petty L, Petty A, Lowther E, Yu Y, 23 and Me Research Team, Highland H, Avery C, Harris KM, Gordon R, Beilby J, Viljoen K, Jones R, Huff C, Kraft SJ. 2023. Discovery of 36 loci significantly associated with stuttering. Res Sq.

12. Bishop M, SheikhBahaei S. 2023. Brainstem astrocytes regulate breathing and may affect arousal state in rats. bioRxiv.

13. Bloodstein O, Ratner NB, Brundage SB. 2021. A handbook on stuttering. Plural Publishing.

14. Bovolenta P, Liem RK, Mason CA. 1984. Development of cerebellar astroglia: transitions in form and cytoskeletal content. Dev Biol 102:248–259.

15. Bradl M, Lassmann H. 2010. Oligodendrocytes: biology and pathology. Acta Neuropathol 119:37–53.

16. Burman KJ, Bakola S, Richardson KE, Reser DH, Rosa MGP. 2014. Patterns of afferent input to the caudal and rostral areas of the dorsal premotor cortex (6DC and 6DR) in the marmoset monkey. J Comp Neurol 522:3683–3716.

17. Chang S-E, Guenther FH. 2019. Involvement of the Cortico-Basal Ganglia-Thalamocortical Loop in Developmental Stuttering. Front Psychol 10:3088.

18. Chang S-E, Horwitz B, Ostuni J, Reynolds R, Ludlow CL. 2011. Evidence of left inferior frontal-premotor structural and functional connectivity deficits in adults who stutter. Cereb Cortex 21:2507–2518.

19. Chow HM, Garnett EO, Li H, Etchell A, Sepulcre J, Drayna D, Chugani D, Chang S-E. 2020. Linking Lysosomal Enzyme Targeting Genes and Energy Metabolism with Altered Gray Matter Volume in Children with Persistent Stuttering. Neurobiol Lang (Camb) 1:365–380.

20. Craig-McQuaide A, Akram H, Zrinzo L, Tripoliti E. 2014. A review of brain circuitries involved in stuttering. Front Hum Neurosci 8:884.

21. Cravedi E, Deniau E, Giannitelli M, Xavier J, Hartmann A, Cohen D. 2017. Tourette syndrome and other neurodevelopmental disorders: a comprehensive review. Child Adolesc Psychiatry Ment Health 11:59.

22. Dahl D, Rueger DC, Bignami A, Weber K, Osborn M. 1981. Vimentin, the 57 000 molecular weight protein of fibroblast filaments, is the major cytoskeletal component in immature glia. Eur J Cell Biol 24:191–196.

23. Debus E, Weber K, Osborn M. 1983. Monoclonal antibodies to desmin, the muscle-specific intermediate filament protein. EMBO J 2:2305–2312.

24. Del Pino I, Rico B, Marín O. 2018. Neural circuit dysfunction in mouse models of neurodevelopmental disorders. Curr Opin Neurobiol 48:174–182.

25. Diamantaki M, Frey M, Berens P, Preston-Ferrer P, Burgalossi A. 2016. Sparse activity of identified dentate granule cells during spatial exploration. eLife 5.

26. Dominy SS, Lynch C, Ermini F, Benedyk M, Marczyk A, Konradi A, Nguyen M, Haditsch U, Raha D, Griffin C, Holsinger LJ, Arastu-Kapur S, Kaba S, Lee A, Ryder MI, Potempa B, Mydel P, Hellvard A, Adamowicz K, Hasturk H, Walker GD, Reynolds EC, Faull RLM, Curtis MA, Dragunow M, Potempa J. 2019. Porphyromonas gingivalis in Alzheimer’s disease brains: Evidence for disease causation and treatment with small-molecule inhibitors. Sci Adv 5:eaau3333.

27. Donaher J, Richels C. 2012. Traits of attention deficit/hyperactivity disorder in school-age children who stutter. J Fluency Disord 37:242–252.

28. Eng LF, Ghirnikar RS, Lee YL. 2000. Glial fibrillary acidic protein: GFAP-thirty-one years (1969-2000). Neurochem Res 25:1439–1451.

29. Forny-Germano L, Lyra e Silva NM, Batista AF, Brito-Moreira J, Gralle M, Boehnke SE, Coe BC, Lablans A, Marques SA, Martinez AMB, Klein WL, Houzel J-C, Ferreira ST, Munoz DP, De Felice FG. 2014. Alzheimer’s disease-like pathology induced by amyloid-β oligomers in nonhuman primates. J Neurosci 34:13629–13643.

30. Frigerio-Domingues CE, Gkalitsiou Z, Zezinka A, Sainz E, Gutierrez J, Byrd C, Webster R, Drayna D. 2019. Genetic factors and therapy outcomes in persistent developmental stuttering. J Commun Disord 80:11–17.

31. Frigerio Domingues CE, Raza MH, Han T-U, Barnes T, Shaw P, Sudre G, Riazuddin S, Morell RJ, Drayna D. 2022. Mutations in *ZBTB20* in individuals with persistent stuttering. medRxiv.

32. Garnett EO, Chow HM, Nieto-Castañón A, Tourville JA, Guenther FH, Chang S-E. 2018. Anomalous morphology in left hemisphere motor and premotor cortex of children who stutter. Brain 141:2670–2684.

33. Gilmore JH, Lin W, Corouge I, Vetsa YSK, Smith JK, Kang C, Gu H, Hamer RM, Lieberman JA, Gerig G. 2007. Early postnatal development of corpus callosum and corticospinal white matter assessed with quantitative tractography. AJNR Am J Neuroradiol 28:1789–1795.

34. Gordon GRJ, Choi HB, Rungta RL, Ellis-Davies GCR, MacVicar BA. 2008. Brain metabolism dictates the polarity of astrocyte control over arterioles. Nature 456:745– 749.

35. Guillamon-Vivancos T, Tyler WA, Medalla M, Chang WW-E, Okamoto M, Haydar TF, Luebke JI. 2019. Distinct Neocortical Progenitor Lineages Fine-tune Neuronal Diversity in a Layer-specific Manner. Cereb Cortex 29:1121–1138.

36. Halassa MM, Fellin T, Haydon PG. 2009. Tripartite synapses: roles for astrocytic purines in the control of synaptic physiology and behavior. Neuropharmacology 57:343–346.

37. Han T-U, Root J, Reyes LD, Huchinson EB, Hoffmann J du, Lee W-S, Barnes TD, Drayna D. 2019. Human GNPTAB stuttering mutations engineered into mice cause vocalization deficits and astrocyte pathology in the corpus callosum. Proc Natl Acad Sci USA 116:17515–17524.

38. Höbler F, Bitan T, Tremblay L, De Nil L. 2022. Differences in implicit motor learning between adults who do and do not stutter. Neuropsychologia 174:108342.

39. Iadecola C, Nedergaard M. 2007. Glial regulation of the cerebral microvasculature. Nat Neurosci 10:1369–1376.

40. Jarvis ED. 2019. Evolution of vocal learning and spoken language. Science 366:50–54.

41. Jürgens U. 2002. Neural pathways underlying vocal control. Neurosci Biobehav Rev 26:235–258.

42. Kang C, Riazuddin S, Mundorff J, Krasnewich D, Friedman P, Mullikin JC, Drayna D. 2010. Mutations in the lysosomal enzyme-targeting pathway and persistent stuttering. N Engl J Med 362:677–685.

43. Kazemi N, Estiar MA, Fazilaty H, Sakhinia E. 2018. Variants in GNPTAB, GNPTG and NAGPA genes are associated with stutterers. Gene 647:93–100.

44. Kettenmann H, Hanisch U-K, Noda M, Verkhratsky A. 2011. Physiology of microglia. Physiol Rev 91:461–553.

45. Key G, Petersen JL, Becker MH, Duchrow M, Schlüter C, Askaa J, Gerdes J. 1993. New antiserum against Ki-67 antigen suitable for double immunostaining of paraffin wax sections. J Clin Pathol 46:1080–1084.

46. Khakh BS, Sofroniew MV. 2015. Diversity of astrocyte functions and phenotypes in neural circuits. Nat Neurosci 18:942–952.

47. Kim KS, Daliri A, Flanagan JR, Max L. 2020. Dissociated Development of Speech and Limb Sensorimotor Learning in Stuttering: Speech Auditory-motor Learning is Impaired in Both Children and Adults Who Stutter. Neuroscience 451:1–21.

48. Kraaimaat FW, Vanryckeghem M, Van Dam-Baggen R. 2002. Stuttering and social anxiety. J Fluency Disord 27:319–30; quiz 330.

49. Krämer-Albers E-M, Werner HB. 2023. Mechanisms of axonal support by oligodendrocyte-derived extracellular vesicles. Nat Rev Neurosci 24:474–486.

50. Krol A, Wimmer RD, Halassa MM, Feng G. 2018. Thalamic reticular dysfunction as a circuit endophenotype in neurodevelopmental disorders. Neuron 98:282–295.

51. Kuhn S, Gritti L, Crooks D, Dombrowski Y. 2019. Oligodendrocytes in development, myelin generation and beyond. Cells 8.

52. Lu C, Peng D, Chen C, Ning N, Ding G, Li K, Yang Y, Lin C. 2010. Altered effective connectivity and anomalous anatomy in the basal ganglia-thalamocortical circuit of stuttering speakers. Cortex 46:49–67.

53. Maguire GA, Nguyen DL, Simonson KC, Kurz TL. 2020. The pharmacologic treatment of stuttering and its neuropharmacologic basis. Front Neurosci 14:158.

54. Maguire GA, Yoo BR, SheikhBahaei S. 2021. Investigation of risperidone treatment associated with enhanced brain activity in patients who stutter. Front Neurosci 15:598949.

55. Marina N, Turovsky E, Christie IN, Hosford PS, Hadjihambi A, Korsak A, Ang R, Mastitskaya S, Sheikhbahaei S, Theparambil SM, Gourine AV. 2018. Brain metabolic sensing and metabolic signaling at the level of an astrocyte. Glia 66:1185–1199.

56. Meyer N, Richter N, Fan Z, Siemonsmeier G, Pivneva T, Jordan P, Steinhäuser C, Semtner M, Nolte C, Kettenmann H. 2018. Oligodendrocytes in the mouse corpus callosum maintain axonal function by delivery of glucose. Cell Rep 22:2383–2394.

57. Miller CT, Thomas AW, Nummela SU, de la Mothe LA. 2015. Responses of primate frontal cortex neurons during natural vocal communication. J Neurophysiol 114:1158– 1171.

58. Molina-Gonzalez I, Holloway RK, Jiwaji Z, Dando O, Kent SA, Emelianova K, Lloyd AF, Forbes LH, Mahmood A, Skripuletz T, Gudi V, Febery JA, Johnson JA, Fowler JH, Kuhlmann T, Williams A, Chandran S, Stangel M, Howden AJM, Hardingham GE, Miron VE. 2023. Astrocyte-oligodendrocyte interaction regulates central nervous system regeneration. Nat Commun 14:3372.

59. Mollaei F, Mersov A, Woodbury M, Jobst C, Cheyne D, De Nil L. 2021. White matter microstructural differences underlying beta oscillations during speech in adults who stutter. Brain Lang 215:104921.

60. Moore S, Meschkat M, Ruhwedel T, Trevisiol A, Tzvetanova ID, Battefeld A, Kusch K, Kole MHP, Strenzke N, Möbius W, de Hoz L, Nave K-A. 2020. A role of oligodendrocytes in information processing. Nat Commun 11:5497.

61. Morquette P, Verdier D, Kadala A, Féthière J, Philippe AG, Robitaille R, Kolta A. 2015. An astrocyte-dependent mechanism for neuronal rhythmogenesis. Nat Neurosci 18:844–854.

62. Mullen RJ, Buck CR, Smith AM. 1992. NeuN, a neuronal specific nuclear protein in vertebrates. Development 116:201–211.

63. Nimmerjahn A, Kirchhoff F, Helmchen F. 2005. Resting microglial cells are highly dynamic surveillants of brain parenchyma in vivo. Science 308:1314–1318.

64. Paolicelli RC, Bolasco G, Pagani F, Maggi L, Scianni M, Panzanelli P, Giustetto M, Ferreira TA, Guiducci E, Dumas L, Ragozzino D, Gross CT. 2011. Synaptic pruning by microglia is necessary for normal brain development. Science 333:1456–1458.

65. Parkhurst CN, Yang G, Ninan I, Savas JN, Yates JR, Lafaille JJ, Hempstead BL, Littman DR, Gan W-B. 2013. Microglia promote learning-dependent synapse formation through brain-derived neurotrophic factor. Cell 155:1596–1609.

66. Petrides M. 2005. Lateral prefrontal cortex: architectonic and functional organization. Philos Trans R Soc Lond B Biol Sci 360:781–795.

67. Polikowsky HG, Shaw DM, Petty LE, Chen H-H, Pruett DG, Linklater JP, Viljoen KZ, Beilby JM, Highland HM, Levitt B, Avery CL, Mullan Harris K, Jones RM, Below JE, Kraft SJ. 2022. Population-based genetic effects for developmental stuttering. HGG Adv 3:100073.

68. Pool KD, Devous MD, Freeman FJ, Watson BC, Finitzo T. 1991. Regional cerebral blood flow in developmental stutterers. Arch Neurol 48:509–512.

69. Qiu Z, Kala S, Guo J, Xian Q, Zhu J, Zhu T, Hou X, Wong KF, Yang M, Wang H, Sun L. 2020. Targeted Neurostimulation in Mouse Brains with Non-invasive Ultrasound. Cell Rep 32:108033.

70. Reagan LP, Cowan HB, Woodruff JL, Piroli GG, Erichsen JM, Evans AN, Burzynski HE, Maxwell ND, Loyo-Rosado FZ, Macht VA, Grillo CA. 2021. Hippocampal-specific insulin resistance elicits behavioral despair and hippocampal dendritic atrophy. Neurobiol Stress 15:100354.

71. Riaz N, Steinberg S, Ahmad J, Pluzhnikov A, Riazuddin S, Cox NJ, Drayna D. 2005. Genomewide significant linkage to stuttering on chromosome 12. Am J Hum Genet 76:647–651.

72. de Rus Jacquet A, Tancredi JL, Lemire AL, DeSantis MC, Li W-P, O’Shea EK. 2021. The LRRK2 G2019S mutation alters astrocyte-to-neuron communication via extracellular vesicles and induces neuron atrophy in a human iPSC-derived model of Parkinson’s disease. eLife 10.

73. Sahin M, Sur M. 2015. Genes, circuits, and precision therapies for autism and related neurodevelopmental disorders. Science 350.

74. Saur L, Baptista PPA, de Senna PN, Paim MF, do Nascimento P, Ilha J, Bagatini PB, Achaval M, Xavier LL. 2014. Physical exercise increases GFAP expression and induces morphological changes in hippocampal astrocytes. Brain Struct Funct 219:293–302.

75. Schafer DP, Lehrman EK, Kautzman AG, Koyama R, Mardinly AR, Yamasaki R, Ransohoff RM, Greenberg ME, Barres BA, Stevens B. 2012. Microglia sculpt postnatal neural circuits in an activity and complement-dependent manner. Neuron 74:691–705.

76. Schindelin J, Arganda-Carreras I, Frise E, Kaynig V, Longair M, Pietzsch T, Preibisch S, Rueden C, Saalfeld S, Schmid B, Tinevez J-Y, White DJ, Hartenstein V, Eliceiri K, Tomancak P, Cardona A. 2012. Fiji: an open-source platform for biological-image analysis. Nat Methods 9:676–682.

77. Shaw DM, Polikowsky HP, Pruett DG, Chen H-H, Petty LE, Viljoen KZ, Beilby JM, Jones RM, Kraft SJ, Below JE. 2021. Phenome risk classification enables phenotypic imputation and gene discovery in developmental stuttering. Am J Hum Genet 108:2271–2283.

78. SheikhBahaei S, Farhan M, Maguire GA. 2022. Improvement of stuttering after administration of methylphenidate - a case report. Personalized Medicine in Psychiatry 31–32.

79. SheikhBahaei S, Marina N, Rajani V, Kasparov S, Funk GD, Smith JC, Gourine AV. 2023a. Contributions of carotid bodies, retrotrapezoid nucleus neurons and preBötzinger complex astrocytes to the CO2-sensitive drive for breathing. J Physiol (Lond).

80. SheikhBahaei S, Millwater M, Maguire GA. 2023b. Stuttering as a spectrum disorder: A hypothesis. Current Research in Neurobiology:100116.

81. Sheikhbahaei S, Morris B, Collina J, Anjum S, Znati S, Gamarra J, Zhang R, Gourine AV, Smith JC. 2018a. Morphometric analysis of astrocytes in brainstem respiratory regions. J Comp Neurol 526:2032–2047.

82. Sheikhbahaei S, Turovsky EA, Hosford PS, Hadjihambi A, Theparambil SM, Liu B, Marina N, Teschemacher AG, Kasparov S, Smith JC, Gourine AV. 2018b. Astrocytes modulate brainstem respiratory rhythm-generating circuits and determine exercise capacity. Nat Commun 9:370.

83. Sholl DA. 1953. Dendritic organization in the neurons of the visual and motor cortices of the cat. J Anat 87:387–406.

84. Simões CS, Vianney PVR, de Moura MM, Freire MAM, Mello LE, Sameshima K, Araújo JF, Nicolelis MAL, Mello CV, Ribeiro S. 2010. Activation of frontal neocortical areas by vocal production in marmosets. Front Integr Neurosci 4.

85. Simonyan K, Fuertinger S. 2015. Speech networks at rest and in action: interactions between functional brain networks controlling speech production. J Neurophysiol 113:2967–2978.

86. Sofroniew MV, Vinters HV. 2010. Astrocytes: biology and pathology. Acta Neuropathol 119:7–35.

87. Stadelmann C, Timmler S, Barrantes-Freer A, Simons M. 2019. Myelin in the central nervous system: structure, function, and pathology. Physiol Rev 99:1381–1431.

88. Stanfield BB, O’Leary DD, Fricks C. 1982. Selective collateral elimination in early postnatal development restricts cortical distribution of rat pyramidal tract neurones. Nature 298:371–373.

89. Stanfield BB, O’Leary DD. 1985. The transient corticospinal projection from the occipital cortex during the postnatal development of the rat. J Comp Neurol 238:236–248.

90. Theodoni P, Majka P, Reser DH, Wójcik DK, Rosa MGP, Wang X-J. 2020. Structural attributes and principles of the neocortical connectome in the marmoset monkey. BioRxiv.

91. Timsit S, Martinez S, Allinquant B, Peyron F, Puelles L, Zalc B. 1995. Oligodendrocytes originate in a restricted zone of the embryonic ventral neural tube defined by DM-20 mRNA expression. J Neurosci 15:1012–1024.

92. Turk AZ, Bishop M, Adeck A, SheikhBahaei S. 2022. Astrocytic modulation of central pattern generating motor circuits. Glia 70:1506–1519.

93. Turk AZ, Lotfi Marchoubeh M, Fritsch I, Maguire GA, SheikhBahaei S. 2021. Dopamine, vocalization, and astrocytes. Brain Lang 219:104970.

94. Turk AZ, SheikhBahaei S. 2022. Morphometric analysis of astrocytes in vocal production circuits of common marmoset (Callithrix jacchus). J Comp Neurol 530:574–589.

95. Wake H, Moorhouse AJ, Jinno S, Kohsaka S, Nabekura J. 2009. Resting microglia directly monitor the functional state of synapses in vivo and determine the fate of ischemic terminals. J Neurosci 29:3974–3980.

96. Watkins KE, Smith SM, Davis S, Howell P. 2008. Structural and functional abnormalities of the motor system in developmental stuttering. Brain 131:50–59.

97. Yairi E, Ambrose N. 2013. Epidemiology of stuttering: 21st century advances. J Fluency Disord 38:66–87.

